# CPDD: Adapting an autoregressive DNA language model for bidirectional phage genome modeling

**DOI:** 10.64898/2026.11.07.765118

**Authors:** Hao Ding, Nannan Wu, Tianyi Qiu

## Abstract

Autoregressive DNA language models generate sequences efficiently from left to right, but many genome-analysis and sequence-editing problems are naturally conditioned on both flanks. We introduce **CreatiPhage**, a 144.5M-parameter autoregressive DNA language model for bacteriophages, and its diffusion extension, **CreatiPhage Diffusion DNA-LM (CPDD)**. CPDD is a 147.5M-parameter bidirectional discrete diffusion model initialized from the six-epoch CreatiPhage checkpoint. CPDD retains the pretrained Transformer, block attention residuals, and Engram conditional memory, while introducing non-causal attention, a MASK-aware base-5 Engram, centered depthwise convolution, and position-aligned nucleotide readout. Training combines masked and GIDD-style uniformly substituted bases with nucleotide- and amino-acid-level losses over a 10 kb, 32 kb, and whole-source context curriculum. On a screened panel of 522 phage CDSs, CPDD achieves higher random-mask restoration accuracy than a comparable diffusion model and performs well above codon-level references. In a mixed-corruption test, it restores 42.4% of visible incorrect bases. Frozen CPDD representations also support *infA* fitness prediction and seven-class phage virion protein subtype classification. Controlled evaluations further characterize how restoration depends on visible context and identify opportunities to improve longer-range sequence recovery. Together, these results support scaffold-conditioned sequence modeling as a step towards modular design of phage genomes.

## 1 Introduction

DNA language models have expanded the scope of sequence analysis and generation, with autoregressive models such as HyenaDNA [1], Evo [2], and megaDNA [3] supporting long-context nucleotide modeling. Many sequence-restoration and editing tasks, however, begin with an existing genomic scaffold: the surrounding sequence is known, while selected positions or regions require reconstruction. These tasks call for models that use both upstream and downstream context while preserving the observed sequence. For bacteriophage genomes, modeling selected regions within their surrounding sequence is a foundational capability for the longer-term goal of modular genome design.

Bidirectional encoders such as DNABERT [4] and Nucleotide Transformer [5] learn representations from both sides of a sequence position. Discrete diffusion extends bidirectional conditioning to generation through repeated recovery of corrupted tokens [6, 7]. LLaDA [8] demonstrates diffusion language modeling through training from scratch, while DiffuLLaMA [9] shows how pretrained autoregressive models can be adapted to this setting. D3LM [10] and PhaGen [11] further explore diffusion-based modeling of DNA and phage genomes. Together, these studies motivate the use of bidirectional denoising for sequence restoration and raise a practical question: how can an autoregressive phage model be adapted to recover uncertain regions within an observed scaffold?

Here, we introduce CreatiPhage, a 144.5M-parameter autoregressive DNA language model for bacteriophages, and its bidirectional diffusion extension, **CreatiPhage Diffusion DNA-LM (CPDD)** (Fig. 1). CreatiPhage incorporates Engram [12] and Block Attention Residuals (AttnRes) [13] for local sequence memory and learned aggregation across network depth. CPDD is initialized from the six-epoch CreatiPhage checkpoint and expands the model to 147.5M parameters through adaptations for corrupted inputs. The conversion introduces bidirectional attention, position-aligned prediction, and memory operations that accommodate masked bases. Motivated by the mixed corruption used in GIDD [14], training includes both masked positions and uniformly substituted bases, allowing the model to learn from missing information and visible sequence errors. Nucleotide reconstruction is combined with amino-acid-level supervision, and training progresses from short windows to whole-source context.

**Figure 1:**
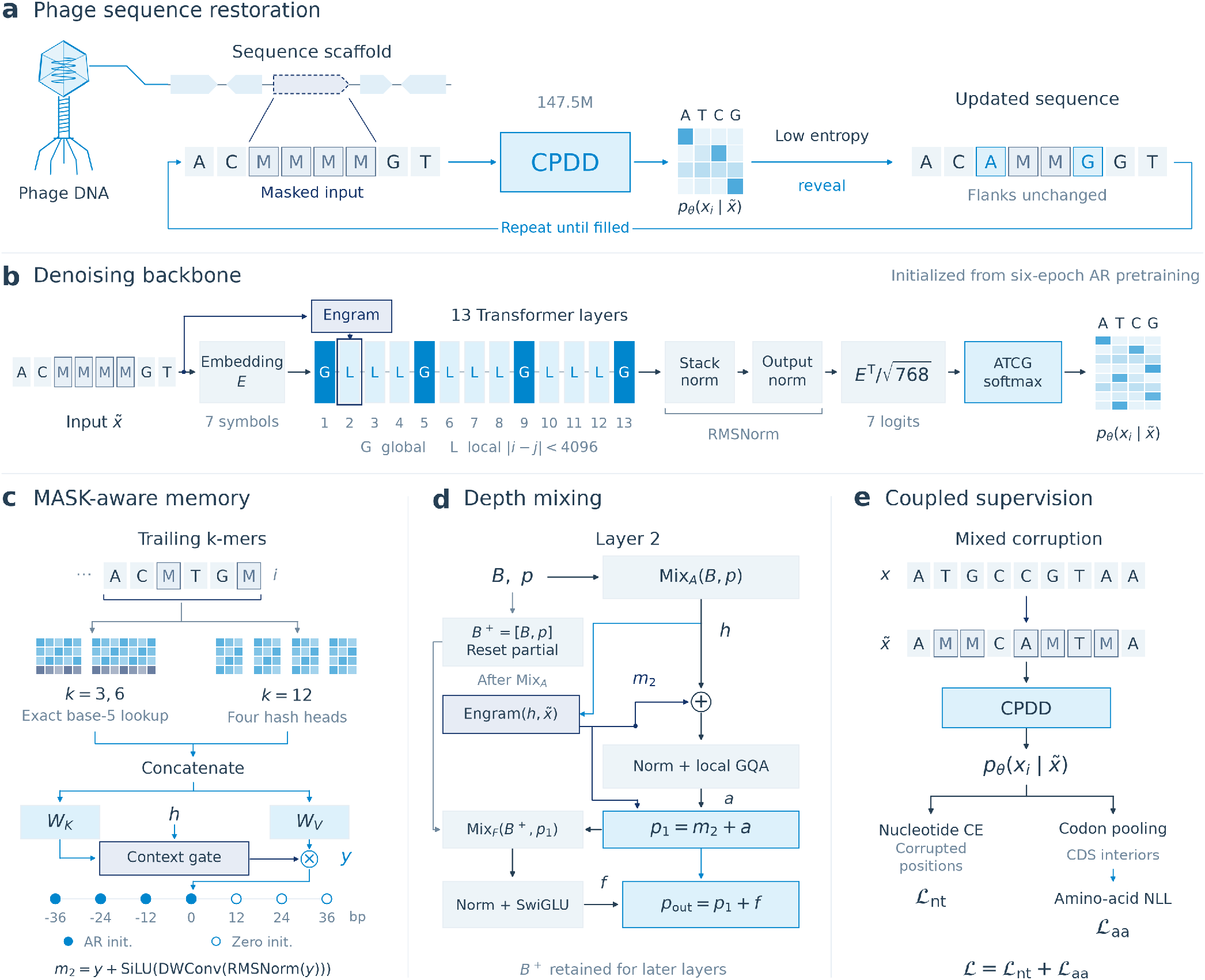
CPDD architecture and sequence restoration. **(a)** One restoration round. The full masked input enters CPDD; the remaining MASK positions with lowest entropy are revealed by argmax or categorical sampling. The updated sequence is fed back until the fixed-length target is filled, preserving visible flanks and earlier writes. Green target bases are newly revealed. All phage, gene, token, memory, and probability graphics are schematic, not experimental outputs. **(b)** The 13-layer backbone initialized from six-epoch AR pretraining. G/L indicate global/local attention within complete Transformer layers, which also contain Block AttnRes and SwiGLU. Corrupted token IDs additionally feed Engram at layer 2. Stack and output RMSNorms precede the scaled, tied seven-logit readout; ATCG selection occurs before softmax. Predictions remain position aligned. Special boundary tokens are omitted from the schematic. **(c)** Trailing *k*-mer lookup uses exact base-5 tables (*k* = 3, 6) and four hash heads (*k* = 12). Ochre table cells indicate new MASK-containing entries. Separate key/value projections supply a gate comparing normalized keys and hidden states *h*. Gated values *y* undergo centered depthwise convolution with dilation 12 and a residual addition. Filled taps inherit AR weights; open taps start at zero and remain trainable. **(d)** Layer 2 at a depth-block boundary. The attention mixer reads the old (*B, p*) before saving *p* in *B*^+^ and resetting the partial accumulator. Engram uses the resulting *h* and corrupted tokens. Its update *m*_2_ enters both the attention input and the new partial *p*_1_. The FFN mixer reads (*B*^+^, *p*_1_); *B*^+^ is retained. Mix denotes Block AttnRes. **(e)** Nucleotide cross-entropy and amino-acid negative log-likelihood share four-base probabilities. Ochre-outlined nucleotide letters are uniform-base draws. Amino-acid supervision pools synonymous codons over eligible whole-CDS interiors after strand reorientation and clamping visible bases; there is no separate protein head.

We first characterize CreatiPhage through sequence-analysis benchmarks and the properties of its generated sequences, then evaluate CPDD through sequence restoration, prediction at visible-error sites, and frozen-representation tasks. CPDD achieves higher random-mask restoration accuracy than a comparable diffusion model on the screened evaluation panel and recovers a fraction of deliberately substituted bases. Controlled tests further examine how recovery depends on the amount and arrangement of visible context, revealing strong local restoration alongside limitations in long-gap recovery and the use of distant sequence information. These results characterize the capabilities and current boundaries of autoregressive-to-diffusion adaptation for scaffold-conditioned phage sequence modeling, providing a basis for further work towards modular design of phage genomes.

## 2 Results

### 2.1 Benchmark performance of the autoregressive base

We first evaluated the autoregressive model that supplies CPDD’s initialization. Benchmark datasets were reconstructed from the sources described by megaDNA [3], and the evaluations were rerun for both models. The six-epoch CreatiPhage AR model (144.5M parameters) and an approximately 145M-parameter reference were assessed across four benchmark families (Fig. 2a–c).

**Figure 2:**
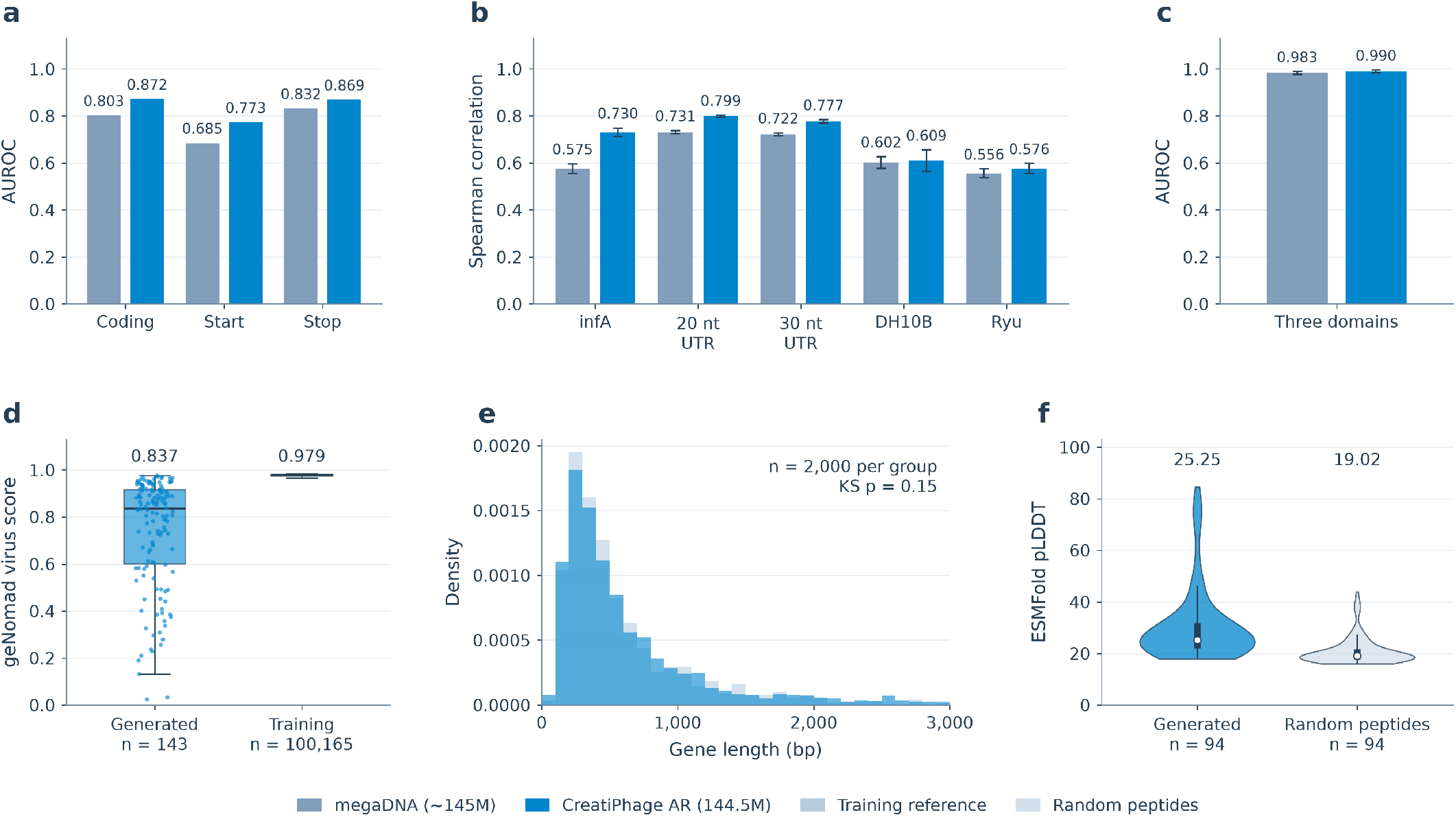
Benchmark performance and generation characteristics of the autoregressive base. **(a–c)** CreatiPhage AR (144.5M parameters) and megaDNA (approximately 145M parameters), evaluated on datasets reconstructed from the sources described by megaDNA. **(a)** Essentiality AUROC for 62 phage *λ* genes, scored by coding-window, startcodon, and stop-codon perturbations. **(b)** Spearman correlations for *infA* fitness, 20 nt and 30 nt UTR libraries, DH10B translation efficiency, and the Ryu dataset (*n* = 4, 599, 11, 692, 11, 889, 1, 050, and 1, 339, respectively). **(c)** Macro one-versus-rest AUROC for bacteria/archaea/phage classification (*n* = 1, 500). Bars in b–c show five-fold means with ±1 standard deviation. **(d)** geNomad virus scores for 143 CreatiPhage AR sequences and 100,165 annotated training sequences or provirus segments. **(e)** Overlaid gene-length density histograms with 100 bp bins and 2,000 genes per group. The displayed range is 0–3,000 bp; densities and the two-sided KS test use the full samples. **(f)** Violin plots of ESMFold pLDDT for 94 generated marker proteins and length-matched random peptides; widths show kernel density estimates, white dots mark medians, and thick bars indicate interquartile ranges. Boxes in d also span the interquartile range, with median lines; points show all generated-sequence scores. Whiskers in d and f extend to the most extreme values within 1.5 times the interquartile range. Numbers above distributions are medians.

For CreatiPhage AR and the size-matched reference, respectively, phage *λ* essentiality AUROCs were 0.872 and 0.803 for coding-window perturbations, 0.773 and 0.685 for start-codon perturbations, and 0.869 and 0.832 for stop-codon perturbations (Fig. 2a). Mean Spearman correlations were 0.730 and 0.575 for *infA* mutational effects; the corresponding translation-related scores were 0.799 and 0.731 for the 20 nt UTR library, 0.777 and 0.722 for the 30 nt library, 0.609 and 0.602 for DH10B, and 0.576 and 0.556 for the Ryu dataset (Fig. 2b). Sequence-domain classification AUROCs were 0.990 and 0.983 (Fig. 2c).

The AR model also generated long sequences with phage-associated features. Across 143 generated sequences, mean length was 47.2 kb and the median geNomad virus score was 0.837 (Fig. 2d). Predicted gene lengths followed a similar distribution to the training reference (two-sample Kolmogorov–Smirnov statistic *D* = 0.036, with 2,000 genes per group; Fig. 2e). Generated marker proteins had a median ESMFold pLDDT of 25.25, compared with 19.02 for length-matched random peptides (94 pairs; Fig. 2f).

These results establish a capable AR starting point for diffusion adaptation.

### 2.2 CPDD restores local sequence and corrects visible errors

We next evaluated scaffold-based restoration and frozen representations, using the public masked-diffusion model PhaGen as a comparable reference (Fig. 3). Mean Spearman correlations on *infA* fitness were 0.736 and 0.698 for CPDD and the reference, respectively; the corresponding macro-F1 scores on seven-class phage virion protein (PVP) subtype classification were 0.809 and 0.715 (Fig. 3a).

**Figure 3:**
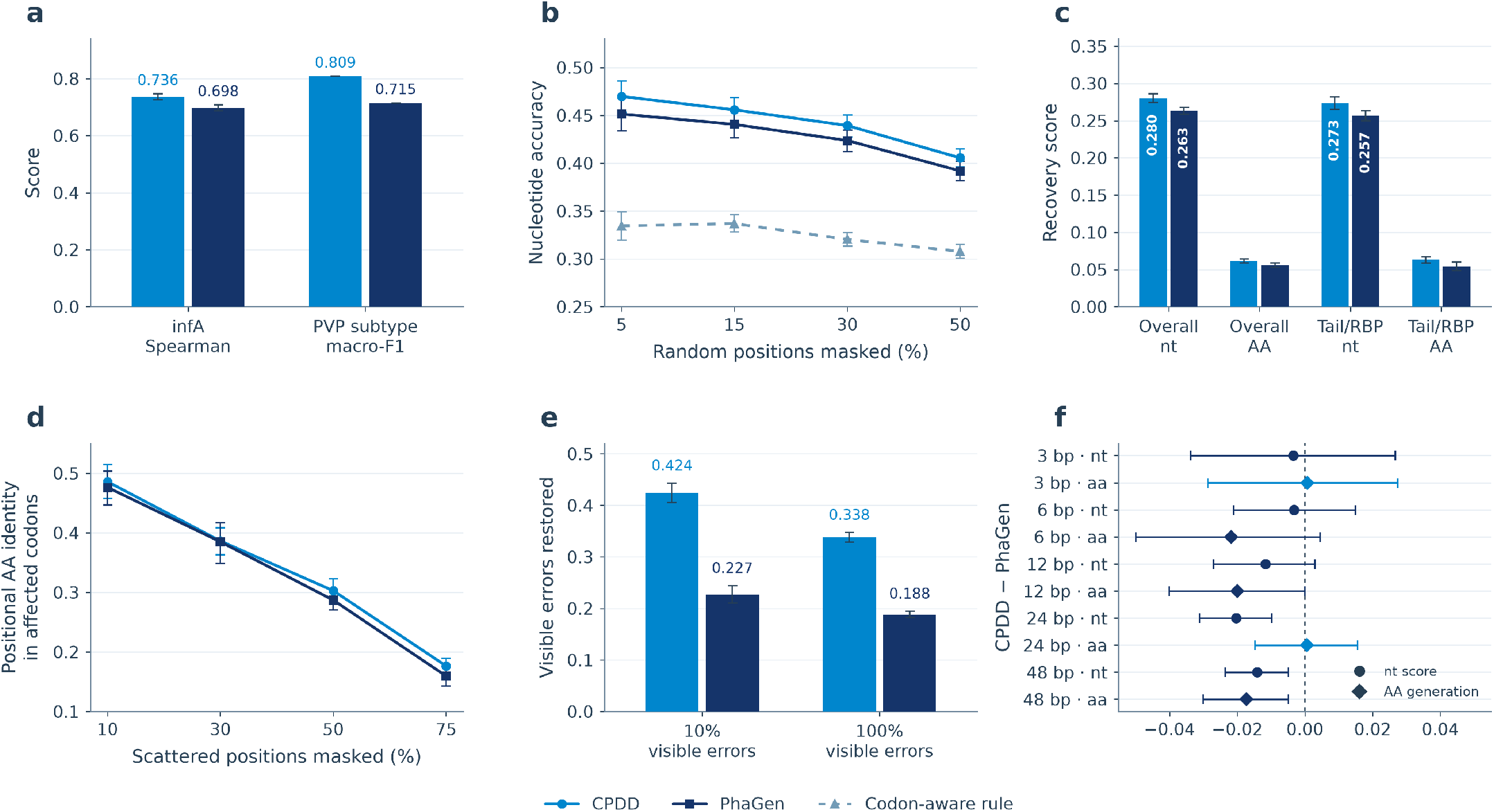
CPDD and PhaGen on scaffold-based restoration tasks. All CPDD results use the final checkpoint. **(a)** Representation benchmarks with frozen embeddings: *infA* fitness (4,600 variants including wild type; mean Spearman correlation over five folds ±1 SD) and seven-class PVP subtype classification (similarity-40 split; macro-F1, single split). **(b)** Nucleotide accuracy for single-pass restoration of randomly masked positions; the dashed line is the codon-aware rule. **(c)** Recovery of 100 functional modules (64-step categorical generation; positional AA identity), overall and for tail-fiber/receptor-binding proteins (*n* = 20). **(d)** Positional AA identity in affected codons after 64-step categorical scattered refill (24 sources). **(e)** Fraction of visible incorrect bases restored to the truth when 30% of positions are corrupted and 10% or 100% of corruptions are visible substitutions. For PhaGen, which was trained with masking only, this is an adapted stress test rather than its native correction interface. **(f)** Paired CPDD− PhaGen differences for contiguous 3–48 bp gaps: nucleotide accuracy from a single forward pass (circles) and positional AA identity after 64-step categorical generation (diamonds). Panels b, e, and f use 522 CDSs in 70 Mash clusters, panel d uses 24 source representatives, and panel c uses 100 modules. Error bars are 95% bootstrap intervals over these units unless noted.

For single-pass restoration of randomly masked positions, accuracies for CPDD and the reference were 0.470 and 0.451 at 5% masking, and 0.406 and 0.392 at 50%, respectively. Both models stayed well above a codon-aware rule that uses only codon-position frequencies and visible bases in the same codon (0.308– 0.337; Fig. 3b). Across 100 functional modules, nucleotide accuracies were 0.280 and 0.263 and amino-acid (AA) identities were 0.062 and 0.056 for CPDD and the reference, respectively; tail-fiber and receptor-binding protein results are shown separately (Fig. 3c). Scattered protein refill showed similar profiles across masking rates (Fig. 3d).

Visible-error prediction was assessed under a mixed-corruption protocol. With 30% of positions corrupted and 10% of those corruptions left as visible substitutions, restoration rates were 42.4% and 22.7% for CPDD and the reference, respectively; when all corruptions were visible, the rates were 33.8% and 18.8% (Fig. 3e). The fractions of visible errors retained unchanged were 20.7% and 52.0% in the first condition. This behavior is consistent with CPDD’s training on substituted as well as masked bases. On contiguous gaps of 3–48 bp, the paired CPDD-minus-reference differences were small and predominantly negative, with magnitudes of 0.003– 0.020 in nucleotide accuracy and up to 0.022 in AA identity (Fig. 3f).

### 2.3 Sequence restoration within a coding scaffold

Figure 4 follows both tasks on a single 753 bp CDS from the 24-source panel. The CDS was chosen as the target whose mean scattered-refill accuracy across 10%, 30%, 50%, and 75% masking was closest to the panel median. This selection did not use the visible-error correction results in panel b; the 5^*′*^ excerpt in panel a was fixed by its position next to the start codon. With 30% of positions masked, 64-step categorical generation restored 35.4% of the 226 masked bases across the CDS. The displayed 5^*′*^ excerpt lies next to the start codon and is locally easier (10 of 12 masked bases restored; Fig. 4a).

**Figure 4:**
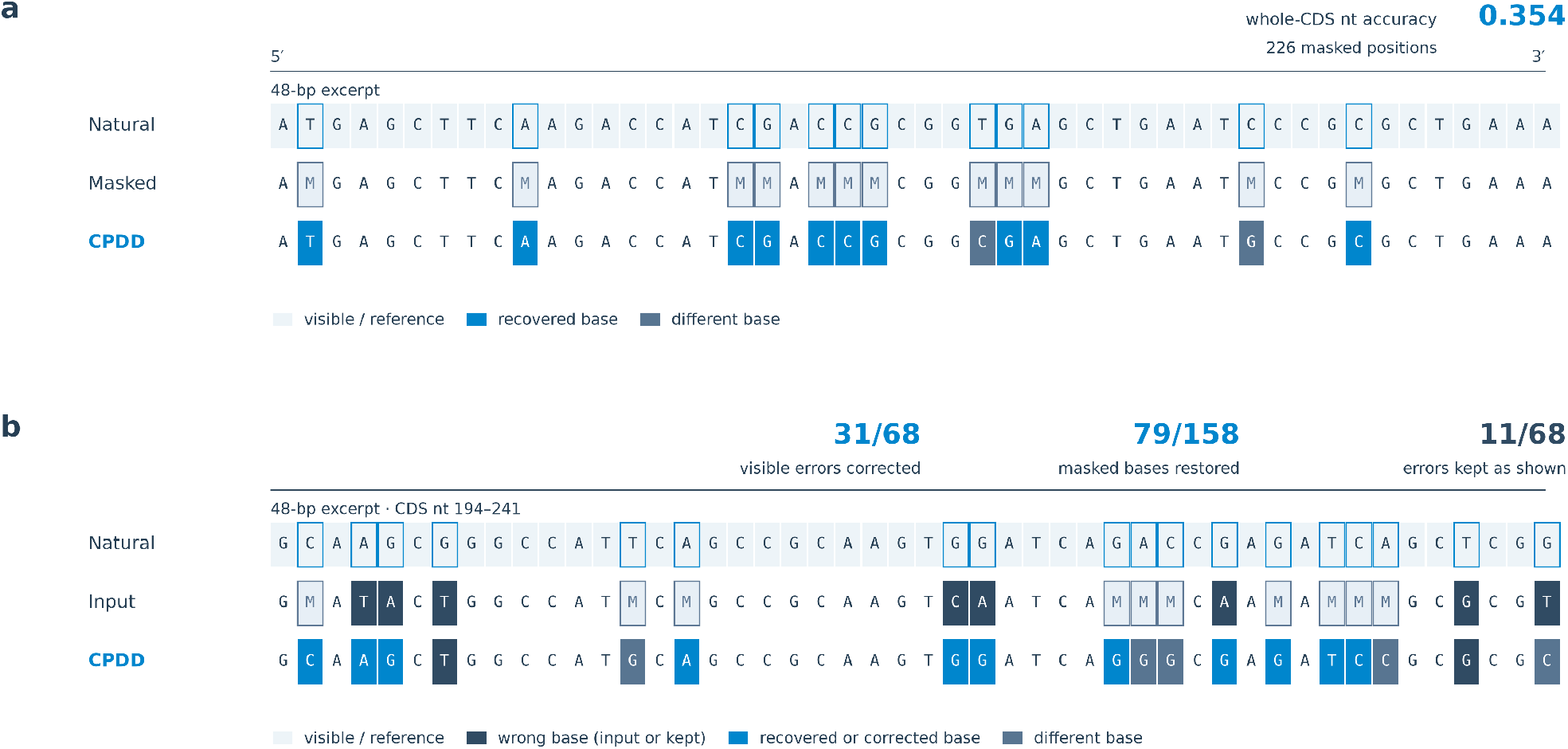
Restoration and error correction on one CDS. The 753 bp CDS was selected by a fixed median rule; all outputs are from the final CPDD checkpoint. **(a)** Masked-base restoration with 30% of positions masked and 64-step categorical generation. Rows show the natural sequence, the masked input (M), and the CPDD output for the 48 bp 5^*′*^ excerpt; outlined bases were masked. The value at top right is nucleotide accuracy over all 226 masked positions. **(b)** Single-pass correction with 30% of positions corrupted: 68 visible incorrect bases (dark in the input) and 158 masked positions. The excerpt is the 48 bp window with the most visible errors (CDS nt 194–241). In the output, bright blue marks restored or corrected bases, mid-blue other bases, and dark tiles visible errors left unchanged. Counts at top are for the whole CDS. Per-position outputs reproduce the cached evaluation job exactly.

The same CDS was then corrupted at 30% of positions, with 68 incorrect bases left visible and 158 positions masked. In a single forward pass, CPDD returned 31 of the 68 visible errors to the correct base and left only 11 unchanged, while restoring 79 of the 158 masked positions. In the 48 bp window with the most visible errors, five of eight errors were corrected (Fig. 4b). Thus, given a scaffold, CPDD both fills masked positions and treats visible bases as evidence rather than fixed truth.

### 2.4 Long gaps approach codon-usage statistics

Random masking leaves visible bases interspersed within a CDS, whereas a continuous gap removes these local clues. We therefore asked whether recovery within longer masked regions reflects sequence-specific information or mainly base and codon preferences. A model-free reference ladder provides a descriptive scale (Fig. 5a; Methods, Section 3.14). With whole CDSs masked, rules that use only base or codon-position frequencies reach 0.264–0.372 nucleotide accuracy. Knowing the encoded protein and choosing synonymous codons would reach 0.771–0.868. CPDD reaches 0.361 and PhaGen 0.312, within the range of the simpler statistics; this level of nucleotide agreement alone does not establish recovery of the original gene.

**Figure 5:**
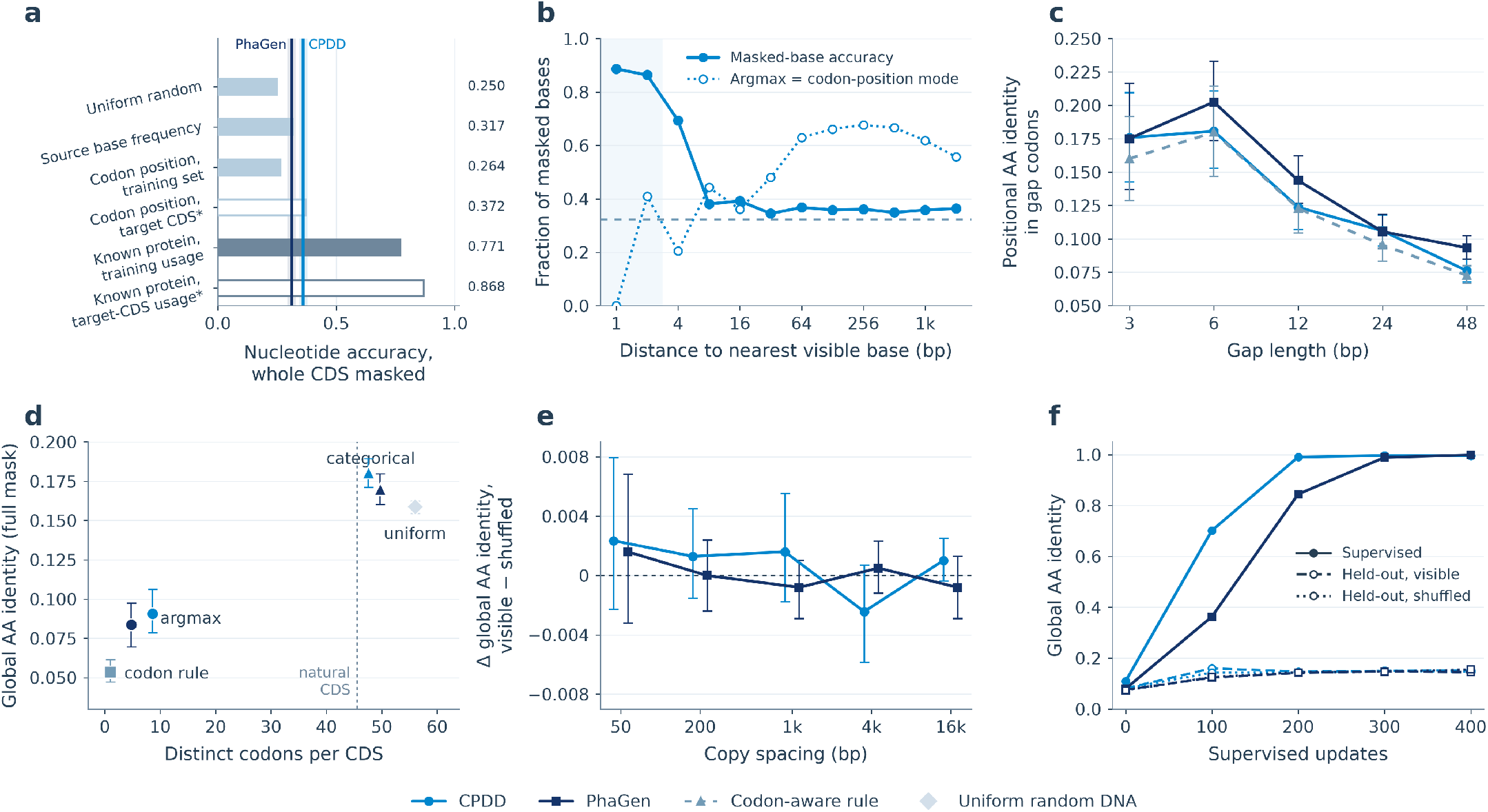
Long-gap behavior of CPDD and PhaGen. **(a)** Model-free reference ladder for whole masked CDSs on the 24-source panel (17,238 coding bases). Light bars use base or codon-position frequencies; dark bars use the true protein with training or target-CDS codon usage. Open bars marked with an asterisk use the evaluated CDS itself to estimate codon-position frequencies or codon usage; the source-base reference uses the complete clean source, including the target. Reference bars average per-CDS accuracy equally across the 24 sources. Vertical lines show source-mean single-pass argmax accuracy for CPDD and PhaGen with whole CDSs masked, with shaded 95% intervals. Algorithms, information access, and aggregation are specified in Methods, Section 3.14. **(b)** Full-mask scoring on the 22-CDS panel, binned by distance to the nearest visible base: masked-base accuracy and the fraction of argmax calls equal to the codon-position mode. The shaded bins are dominated by start/stop codons; the dashed line is the codon-aware rule. **(c)** Positional AA identity of generated gap codons (522 CDSs in 70 clusters, 64 categorical steps) for CPDD, PhaGen, and the codon-aware rule. **(d)** Full-mask outputs on the 24-source panel: distinct codons per CDS versus global-alignment AA identity for one-step argmax (circles) and categorical sampling (triangles; CPDD 32 steps, PhaGen 256 steps), with uniform random DNA and the codon-aware rule. The dotted line marks the natural codon count. **(e)** Paired global-alignment AA-identity differences between a visible verbatim copy and a shuffled copy, inserted 50 bp to 16 kb from the masked original (32 CDSs). Absolute identities across models and spacings range from approximately 0.078 to 0.098. **(f)** Training on copy-bearing contexts (50 bp spacing, eight supervised and eight held-out contexts). Solid lines show supervised contexts; dashed and dotted lines show held-out contexts with visible and shuffled copies. Panels a–e use the final CPDD checkpoint and, where applicable, the released PhaGen weights. Panel f starts from these weights and applies the indicated supervised updates. Error bars are 95% bootstrap intervals over clusters (c) or sources (a, d, e).

To examine where local context contributes, we grouped predictions within fully masked CDSs by distance to the nearest visible base. Accuracy is high next to visible bases but falls to approximately 0.35–0.37 beyond about 32 bp, close to the 0.324 of the codon-aware rule. The argmax also increasingly matches the most frequent base at each codon position (Fig. 5b), consistent with a greater contribution from sequence statistics in the masked interior. Gap infilling on the larger 522-CDS panel shows the same trend: for 24 and 48 bp gaps, AA identity is 0.106 and 0.076 for CPDD and 0.105 and 0.093 for PhaGen, compared with 0.095 and 0.073 for the codon-aware rule (Fig. 5c).

We next examined whether repetitive full-mask outputs depended on decoding (Fig. 5d). One-step argmax repeats a few codons: 8.6 distinct codons per CDS for CPDD and 4.8 for PhaGen, against 45.5 in natural sequences. Multistep categorical sampling restores codon diversity, with AA identities of 0.180 for CPDD and 0.170 for PhaGen, compared with 0.159 for uniform random DNA. Thus, more diverse outputs need not imply accurate recovery of the original sequence. Because these settings differ in both sampling rule and iteration budget, they characterize decoding behavior rather than isolate the effect of sampling alone.

### 2.5 Reading sequence from distant context

A fully masked CDS may not be uniquely determined by its flanking sequence. To test whether recovery also suffers when the required sequence information is available, we placed a verbatim copy of the masked CDS elsewhere in the same input. Each copy-bearing input was paired with a control in which the copy was shuffled, preserving its length and base composition while disrupting its sequence order. This comparison asks whether an intact copy improves recovery beyond composition alone.

Without additional training, single-pass predictions on 32 CDSs showed no consistent recovery advantage from the intact copy across spacings of 50 bp to 16 kb; AA identities across the two models ranged from approximately 0.078 to 0.098 (Fig. 5e). We then tested whether brief task-specific training could improve copy use. Both models received 400 updates on eight copy-bearing contexts at 50 bp spacing, with eight additional contexts held out from these updates. These diagnostic examples came from the original training pool; the holdout applies to this adaptation probe.

Both models learned the eight supervised examples, reaching AA identities of 0.995 for CPDD and 0.999 for PhaGen. Held-out recovery remained around 0.15, with similar performance for intact and shuffled copies (Fig. 5f). For CPDD, the final residue-weighted paired difference was 0.005 (95% interval −0.014 to 0.019). Fitting the supervised examples is compatible with memorization and does not by itself demonstrate a transferable copying rule. Under this small-sample protocol, adaptation therefore showed sequence-specific learning but limited transfer of copy-based recovery to held-out contexts.

## 3 Methods

### 3.1 Sequence collection and coding targets

We reconstructed the megaDNA [3] sequence collection from its NCBI GenBank, Metagenomic Gut Virus (MGV v1.0) [15], and Gut Phage Database (GPD) [16] sources. The collection contains 99,429 source records totaling 4,771,564,509 bp: 16,489 from GenBank, 52,872 from MGV, and 30,068 from GPD. These records include complete genomes, metagenomic contigs, and provirus-derived intervals. We therefore refer to them as source records throughout. The historical autoregressive (AR) split comprises 94,469 training and 4,960 validation records; AR pretraining used the forward strand without reverse-complement augmentation.

For diffusion training, we grouped exact and reverse-complement duplicates and linked source accessions before assigning records to pools. Training records came from the original AR training split; validation and test records came from its validation split, after excluding source groups linked to training. Records containing non-ACGT symbols were excluded. This procedure isolates exact sequence and source overlap, but does not establish general homology separation or erase historical validation exposure.

Coding sequences (CDSs) were obtained from paired geNomad [17] TSV/FAA annotations. We retained complete CDS predictions only when identifiers, coordinates, strand, genetic code, and translated protein agreed. Genetic codes 4, 11, and 15 were supported. CDSs of at most 3,000 bp contributed a whole-CDS target; longer CDSs contributed one deterministically selected, codon-aligned 3,000 bp window. The resulting index contains 5,875,538 training targets (5,794,918 whole CDSs and 80,620 partial windows), 155,178 validation targets, and 154,684 test targets. Training also applies a frozen source-exclusion list. Coordinates are zero-based and half-open; input DNA retains the original source orientation, including for negative-strand targets. The longest eligible source is 95,998 bp, within the model’s 98,304-token limit after adding BOS and EOS.

The offline index stores clean references and target coordinates. Corruption is generated online. Sampling cycles through shuffled source records and draws a primary target within each source, so a source pass does not imply that every target in the 5.88M-target training index has been visited.

### 3.2 Autoregressive backbone and initialization

The AR base has 144,498,102 parameters in a flat, nucleotide-resolution stack of 13 Transformer layers. It uses hidden width 768, 12 query heads, two key/value heads, a SwiGLU feed-forward width of 3,776, QK normalization, rotary position embeddings, and tied input/output embeddings. Layers 1, 5, 9, and 13 use global grouped-query attention; the remaining layers use causal sliding-window attention with window 4,096.

We adapt Engram [12], the conditional-memory architecture developed by DeepSeek-AI and Peking University, to nucleotide sequences, retaining trailing *k*-mer retrieval, hidden-state-dependent gating, and convolutional memory refinement. The module is placed at layer 2. DNA uses canonical nucleotide IDs directly, so the original text-tokenizer compression step is omitted. The *k* = 3 and *k* = 6 branches replace hashed lookup with collision-free base-4 tables, while the *k* = 12 branch uses four hashed heads. In the AR model, memory refinement uses a four-tap causal depthwise convolution with dilation 12. Let **e**_*i*_ denote the concatenated lookup ectors at position *i*, and **h**_*i*_ the incoming hidden state. The retrieved memory is fused with the hidden state through

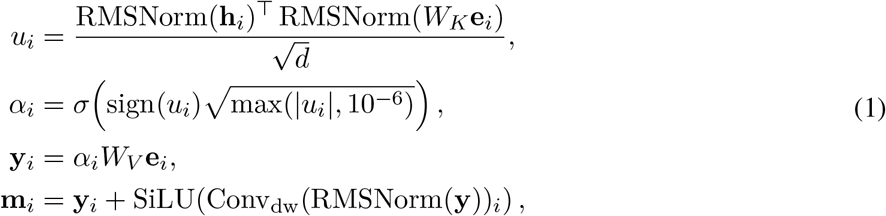

where *d* = 768 and affine projection biases are implicit. The memory update **m**_*i*_ enters both the attention input and the current partial depth block, as detailed below.

We use the Kimi team’s Block AttnRes [13] from Moonshot AI to supply the inputs to the attention and feed-forward sublayers by mixing depth-block representations. The learned pseudo-query, RMS-normalized keys, and softmax weighting over depth follow the original formulation. For sublayer *l* and available block set ℬ_*l*_, the token-wise mixture is

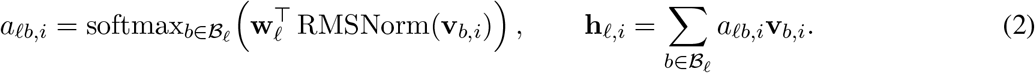

The configured block size is four attention/FFN sublayers. At stack entry, both the initial history element and the partial block equal the token embedding. At each even-numbered layer, the attention-input mixture is computed first; the incoming partial block is then appended to the history and the current accumulator is reset. The history is not truncated. Our implementation evaluates the same depth mixture in two passes over the source states, avoiding stacked hidden-state tensors and accumulating the weighted sum in FP32. To integrate Engram, its gated memory update is added to both the attention input and the partial depth block; the AttnRes weighting rule itself is unchanged.

AR pretraining ran for six epochs (23,274 updates) with a maximum context of 98,304 tokens and approximately 1.57M tokens per update. A 2,000-update warmup preceded cosine decay. Matrix-shaped hidden weights used Muon with peak learning rate 0.02; embedding, normalization, head, and scalar groups used AdamW with peak learning rate 3 × 10^*−*4^ and *β* = (0.9, 0.95). Weight decay was 0.1, the gradient clipping threshold was 2.0, and the minimum learning rate was 10^*−*8^. Computation used FP32 master weights with BF16 autocast.

### 3.3 Conversion to a bidirectional denoiser

CPDD was initialized from this AR checkpoint, with all parameters trainable. The vocabulary adds MASK to the original six rows (BOS/PAD, A, T, C, G, EOS); its embedding is initialized to the mean of the four nucleotide embeddings. Together with the memory expansion below, the conversion gives 147,468,214 parameters.

All attention layers become non-causal (Fig. 1b). The four global layers attend across the input window. The nine local layers use | *i*−*j* | *<* 4096, allowing an interior position to attend to itself and up to 4,095 positions on either side. Global attention uses grouped-query SDPA; local attention uses FlexAttention block masks without constructing a dense token-square mask. After the last layer, two successive learned RMSNorms—the stack norm and the output norm—precede the tied embedding projection. For an embedding matrix *E* ∈ℝ^768*×*7^ whose columns hold the token embeddings, and final partial state **p**_13,*i*_,

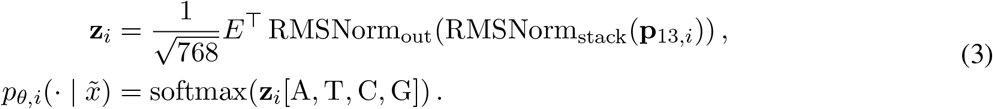

Here *p*_*θ,i*_ is the four-base distribution at position *i*. The readout is aligned to the corrupted position. There is no next-token shift, explicit diffusion-time embedding, or auxiliary protein head.

The exact Engram tables are extended from base-4 to base-5 indexing, with MASK as the fifth digit (Fig. 1c). ATCG-only entries retain their pretrained values; MASK-containing entries are initialized from *N*(0, 0.02^2^). ATCG, MASK-containing, and out-of-vocabulary entries are stored separately so that lookup gradients update only the selected group. The *k* = 12 branch retains its pretrained hash tables and hashes MASK-containing input instead of suppressing it. The *k*-mers themselves remain trailing windows ending at the current position.

Bidirectional memory refinement is supplied by a centered, seven-tap depthwise convolution with dilation 12 (offsets −36, −24, −12, 0, 12, 24, 36 bp): the four inherited causal taps initialize the left/current positions, and the three right-context taps start at zero. AttnRes and its depth-history policy are retained.

Figure 1d makes this ordering explicit for layer 2. With incoming state (*B, p*), the computation is

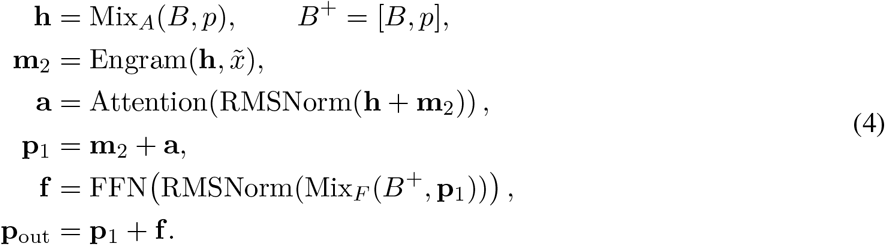

Each other layer has its own attention and FFN depth mixers, with Engram omitted and the partial block carried across non-boundary layers.

### 3.4 Window selection and mixed corruption

A training example begins with a crop centered on the primary target, or the complete source in a whole-source update. In the refined short-window and subsequent stages, additional targets are visited in random order and selected subject to a maximum of eight targets and 300 bp separation between their unexpanded boundaries. Selection stops once the expected corrupted-base budget has been reached. From stage B onward this budget is min(7000, ⌊0.25*L*_*w*_⌋) for a window of *L*_*w*_ bases. It is a selection threshold, not a hard cap on the realized number of supervised bases: the last selected target, stochastic corruption, and boundary expansion can cause an overshoot.

Each target interval may expand by independently sampled 0–100 bp flanks, clipped to the crop and neighboring-target midpoints. In the full branch, half of the interval draws instead retain the exact target boundaries. The partial branch uses expanded intervals. Thus the 300 bp spacing rule applies before expansion, not to the final visible gap.

The full branch selects every position in each supervised interval for corruption. The partial branch draws one rate *r* ∼*U* (0.05, 1) per update and selects positions independently at that rate, ensuring at least one corrupted and one visible base when the interval permits. Let *C*_*j*_ denote the selected positions in interval *j* and *C* = ⋃ _*j*_ *C*_*j*_. Stage A uses pure masking. From stage B onward, a share *γ* ∼ *U* (0, 0.3) is drawn per update. We select a subset *U*_*j*_ ⊆ *C*_*j*_ without replacement, with size min (|*C*_*j*_ |, max(1, round(*γ*|*C*_*j*_ |))) when *γ >* 0 and zero otherwise. Writing *U* =⋃ _*j*_ *U*_*j*_, the corrupted input is

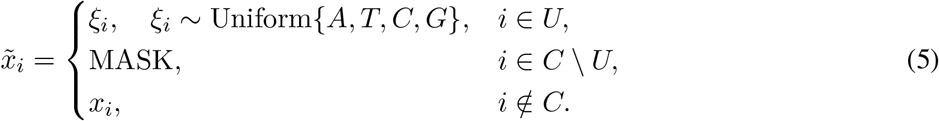

Uniform draws include the true base, so approximately one quarter of substituted positions remain correct. All positions in *C*, including these unchanged draws, are supervised. This borrows the hybrid-noise motivation of GIDD [14]; it does not implement its continuous-time evidence lower bound.

### 3.5 Nucleotide and amino-acid objectives

For one example, nucleotide reconstruction uses cross-entropy over all corrupted positions:

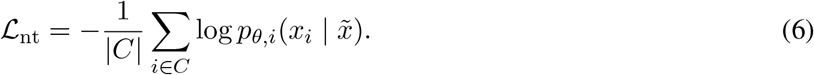

The amino-acid term uses the same four nucleotide probabilities. For each selected whole CDS containing at least one corrupted internal base, we exclude its first and last codons and orient the remaining probabilities in the coding direction. Negative-strand CDSs use a sequence reversal and the class permutation A↔T, C↔G. No input reverse-complement is needed, so targets on both strands share a forward pass. Partial CDS windows do not contribute to this term.

Let *K* collect all internal codons of those eligible CDSs, *i*_*kj*_ denote the position of base *j* in codon *k*, and *τ*_*k*_ the annotated genetic-code mapping. After strand reorientation, define

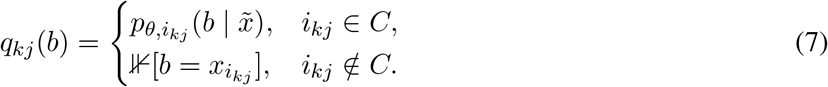

The probability of amino acid *a* and its loss are

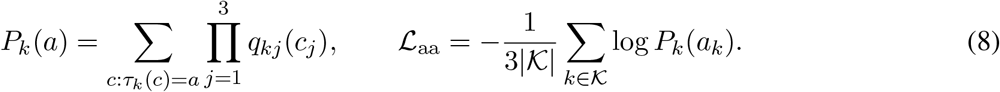

Here *a*_*k*_ is the true residue. The denominator includes all internal codons of each eligible CDS, including fully visible codons, whose loss is zero after clamping. Substituted positions remain predictions even when a draw happens to match the truth. If *K* is empty, the amino-acid loss is zero. The per-example objective is ℒ = ℒ_nt_ + ℒ_aa_; synonymous-codon probabilities are pooled in log space for numerical stability.

The amino-acid term couples the three nucleotide marginals within a codon, but does not define a joint distribution across an entire CDS. For nucleotide cross-entropy alone, an unrestricted predictor is optimized by the conditional base marginals. Independent argmax decoding of weakly informative marginals can therefore yield low-complexity sequences. This observation does not establish the optimum of the combined loss or an information limit for a particular target; those questions require the context controls described below.

### 3.6 Context curriculum and optimization

Diffusion training comprised 46,000 updates across three stages (Table 1). Short-context adaptation began with a 4,096 bp crop radius (at most 8,194 tokens), increasing around update 2,750 to a 5,120 bp radius with a 12% expected-corruption budget and the separated-target rule above. Extended-context adaptation began at update 33,751 and introduced longer crops, occasional whole-source updates, and uniform substitutions. Whole-source training began at update 38,001, retained the same denoiser and objective, and concluded with cosine learningrate decay over updates 40,001–46,000.

**Table 1:** Diffusion training configuration. Updates exclude the preceding AR pretraining. Context lengths include BOS/EOS. “Full” denotes the full-corruption branch, which can contain substituted bases from stage B onward.

| Stage | Diffusion updates | Context | Corruption and schedule |
|---|---|---|---|
| A: short context | 1–33,750 | 8,194, then 10,242 tokens | 50% full; pure MASK; warmup then stable learning rate |
| B: extended context | 33,751–38,000 | up to 32,770 tokens; 15% whole-source updates | 25% full; γ ∼ *U*(0, 0.3); stable learning rate after re-warmup |
| C: whole source | 38,001–46,000 | whole source; model limit 98,304 tokens | 20% full; same hybrid noise; 2,000 stable then 6,000 cosine-decay updates |

Six data-parallel ranks process one source window each per optimizer update. Branch, partial-mask rate, substitution share, and whole-source choice are synchronized; source selection and position/base draws vary across ranks. If rank *s* has *N*_*s*_ corrupted bases and per-example loss ℒ_*s*_, the aggregated gradient corresponds to

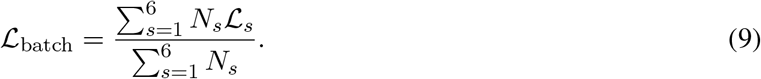

Consequently, the amino-acid term is normalized within each example and then weighted by that example’s corrupted-base count; it is not a separate global codon average.

Diffusion training uses AdamW with *β* = (0.9, 0.95), *ϵ* = 10^*−*8^, weight decay 0.01, and gradient clipping at norm 5.0 after aggregation. Peak learning rates are 10^*−*4^ for new MASK embedding/table rows, 5× 10^*−*5^ for the adaptation group (attention, AttnRes mixers, nucleotide embeddings/readout, Engram projections and convolution), and 2.5×10^*−*5^ for the remaining backbone. This grouping sets learning rates; it does not freeze weights or introduce low-rank adapters. A 500-update warmup precedes the stable phase, with a 300-update re-warmup after context transitions. The final cosine schedule reduced the learning rate from its value at update 40,000 to 10% of each group’s peak at update 46,000. Master weights remain FP32 and forward computation uses BF16 autocast.

### 3.7 Entropy-guided restoration

At inference, the basic sampler masks a target of fixed length *L* and preserves its flanks. For a requested budget *T*, it performs *T* ^*′*^ = min(*T, L*) rounds. At round *k*, the model recomputes the full input and scores the remaining MASK set *M*_*k−*1_ using four-base probabilities (temperature 1 unless specified). It then commits the lowest-entropy positions:

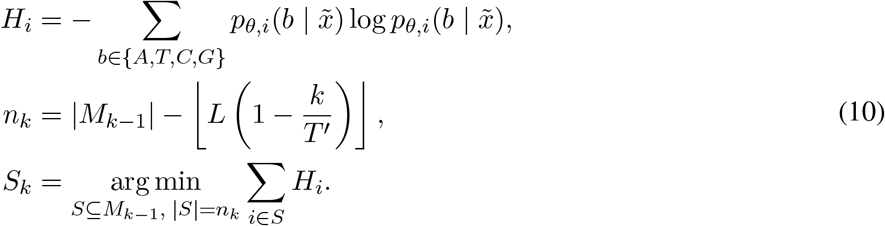

Entropy ties retain sequence order. Values at *S*_*k*_ are selected either by argmax or categorical sampling, and *M*_*k*_ = *M*_*k−*1_ *S*_*k*_. These decoding modes are reported separately. Earlier writes are retained in the basic sampler, and full-context recomputation replaces AR KV caching. The procedure neither changes target length nor predicts an EOS decision within the target.

A separate repair interface scores visible tokens, re-masks low-confidence positions, and refills them. Repair is evaluated using corrected-error and false-edit rates; its availability alone does not establish successful self-correction. When constructing visible-error tests, replacements exclude the true base, unlike the four-base training draws, so every planted error has an unambiguous label.

### 3.8 Autoregressive benchmark evaluation

We reconstructed the benchmark datasets from the sources described by megaDNA [3] and reran the evaluations for both models. The comparison includes the six-epoch CreatiPhage AR checkpoint used to initialize CPDD (144.5M parameters) and the released megaDNA model (approximately 145M parameters). Evaluation covers gene essentiality, mutational effects, translation-related measurements, and sequence-domain classification using the AR weights before diffusion adaptation.

For phage *λ* essentiality, likelihood changes induced by 50 bp coding-window perturbations or start/stopcodon substitutions were scored in whole-genome context. AUROC was computed over 62 labeled genes (34 essential and 28 nonessential). For *infA* mutational effects, a ridge-regression probe used frozen sequence embeddings to predict rich-medium fitness for 4,599 variants. Translation probes used the same regression procedure on random 20 nt and 30 nt 5^*′*^-UTR libraries (*n* = 11, 692 and 11, 889), the top-expression quartile of the DH10B dataset (*n* = 1, 050), and all 1,339 retained records in the reconstructed Ryu translation-efficiency dataset. Sequence-domain classification used a logistic-regression probe on 1,500 fragments (500 each from bacteria, archaea, and phages), reporting macro one-versus-rest AUROC. The probes use each model’s native concatenated representation, whose dimensions differ between architectures.

Regression results are mean Spearman correlations across five shuffled cross-validation folds; classification uses five stratified folds. Error bars show the standard deviation across folds. Essentiality scores are single-dataset AUROCs. Cross-validation partitions individual examples, including variants of one gene for *infA*; pretraining homology overlap has not been comprehensively excluded. We omit the T7 SNP task because its protocol reproduction remained unresolved.

### 3.9 Autoregressive generation evaluation

The AR generation analysis uses 143 sequences longer than 1 kb, sampled from random four-base primers at temperature 0.95 with EOS stopping and a 96 kb length limit. geNomad [17] annotations provide sequence-level virus scores and predicted genes. The training reference contains 100,165 annotated sequence or provirus-segment entries. Gene-length distributions use the existing samples of 2,000 predicted genes from each collection. For protein structure confidence, ESMFold [18] scored all 94 eligible geNomad-marker proteins of at most 1,000 amino acids in the selected marker set, together with 94 length-matched random peptides. These measurements characterize generated sequence features and predicted structure confidence.

### 3.10 Frozen-representation evaluation

Figure 3a follows the public PhaGen evaluation protocols [19], with frozen FP32 embeddings and no gradient updates to either model. CPDD concatenates mean-pooled hidden states from layers 4, 8, and 13 (2,304 dimensions); PhaGen uses its native three-stage mean pooling (964 dimensions), excluding padding from the pooled means. These comparisons use native representations of different dimensions.

For *infA*, the rich-medium fitness measurements of Kelsic et al. [20] provide 4,599 variants and one retained wild-type sequence, each 219 nt long. Five shuffled folds (seed 42) contain 3,680 training and 920 test sequences each. Within each training fold, features are standardized and RidgeCV selects among 500 log-spaced penalties from 10^*−*6^ to 10^6^ by leave-one-out cross-validation. We report mean held-out Spearman correlation and population SD across folds. CPDD pooling includes BOS/EOS tokens; PhaGen receives no added BOS/EOS tokens. The historical AR benchmark in Figure 2 uses 4,599 variants, unstandardized features, and penalties {0.1, 1, 10}, with different fold membership and pooling conventions; values across the two figures should therefore not be interpreted as a before/after effect of diffusion adaptation.

PVP subtype classification uses the PhaVIP-derived data [21] and the official PhaGen similarity-40 split: 7,822 training and 7,079 test CDSs in seven classes. Neither model receives added BOS/EOS tokens in this task. A StandardScaler fitted only on training embeddings precedes class-balanced logistic regression (*C* = 1, L-BFGS, maximum 5,000 iterations, seed 42). Macro-F1 is computed on the fixed test split; it is a single-split result, without a cross-validation error bar.

### 3.11 Diffusion evaluation design and controls

The matched-budget evaluation protocol comprises 24 source representatives screened against the publicly documented CPDD and PhaGen training ranges and specifies 1,660 task–budget combinations per model, including shared 32- and 64-step budgets. The grid covers random masking, scattered refill, 6–96 bp gaps, visible-error detection/correction, and codon diversity; 192 and 768 bp gaps provide longer-gap stress tests. megaDNA is assessed separately on 23 AR continuations, since its continuation interface differs from bidirectional refill. Source representatives, rather than repeated seeds, are the independent units for paired source-level uncertainty estimates.

The final comparison suite contains 522 CDSs grouped into 70 Mash [22] clusters. Targets were screened against the CPDD training data, its AR base, and the public PhaGen training references by accession and reverse-complement matching, Mash distance, local DNA identity (≥95% over ≥100 bp in the target and 512 bp flanks), and protein homology (≥30% identity, ≥80% coverage, *E* ≤ 10^*−*3^); clusters are the statistical units. For this 522-CDS panel, target scores are averaged within each Mash cluster; 95% percentile intervals use 10,000 cluster bootstrap resamples, with model differences paired by cluster. Functional-module recovery uses a separate 100-module panel (Section 3.12). Visible-error restoration is an adapted stress test for PhaGen, which was trained with masking only, rather than its native correction interface. Independence holds within the documented training ranges and frozen thresholds and does not exclude distant homology.

The diagnostic experiments use the final CPDD checkpoint and separate task-specific panels: 22 CDSs for distance-to-context scoring, 24 source representatives for full-mask decoding, and 32 CDSs for visible-copy tests. Controls include model-free nucleotide and amino-acid references, accuracy as a function of distance to visible context, and separate argmax/categorical comparisons. In the latter, a verbatim CDS is placed elsewhere in the same input while the original is masked; shuffling the visible copy preserves its base composition while removing its sequence information. This comparison tests whether available information is used. A separate adaptation probe starts from the final CPDD checkpoint or the released PhaGen weights, performs 400 supervised updates on eight copy-bearing contexts with 50 bp spacing, and evaluates eight held-out contexts with visible or shuffled copies. CPDD uses a fresh optimizer with a peak learning rate of 5 ×10^*−*5^ and a 50-update warmup. Nucleotide recovery, protein identity, internal stops, and codon diversity are interpreted together, without treating any one computational metric as evidence of biological function.

### 3.12 Functional-module evaluation

The panel contains 100 modules from distinct genomes and Mash clusters, with 20 each in tail-fiber/receptor-binding, lysis, lysogeny, packaging/structural, and replication classes. Genes called with Prodigal-gv [17, 23] were assigned functional classes using MMseqs2 [24] matches to PHROG [25] representatives. A module is a run of adjacent, co-oriented CDSs of the same class; a different class or strand ends the run, and a single CDS can form a module. Targets span 300–3,000 bp between outer CDS boundaries, including intervening sequence. For longer modules, a consecutive complete-CDS subset is selected by maximum gene count, then span, then earliest position. The frozen panel contains 99 full modules and one such subset.

The entire target interval is masked, with up to 4,096 bp of visible context on each side, and restored in 64 categorical steps. Nucleotide accuracy covers all masked bases, including intergenic positions. AA identity compares translated CDSs at matching positions in their annotated coding direction, converts valid initial codons to methionine, and removes terminal stop symbols while retaining stop-loss extensions. Matches and denominators are pooled across CDSs within each module, then averaged equally across modules. Confidence intervals resample the 100 modules (or the 20 tail/RBP modules) 10,000 times.

### 3.13 Restoration metrics and decoding

Nucleotide accuracy is the fraction of evaluated bases matching the reference. The scattered-refill and gap AA metrics in Figures 3d,f and F5c compare translations at corresponding codon positions after orienting each CDS in its coding direction. Only codons touched by the mask enter the denominator; unchanged visible codons are excluded. Translation uses the annotated genetic code and retains stop symbols, with no sequence alignment. The module, scattered-refill, and gap results use a budget of up to 64 categorical steps, capped by the number of masked positions, with sampling at temperature 1 with lowest-entropy position selection, retaining original visible bases. Single-pass nucleotide scores instead use argmax predictions.

Figures 5d–f use a different AA metric: global alignment of the translated generated and reference proteins (BLOSUM62; gap opening −11, extension −1). Translation continues through internal stops, and terminal stop symbols are removed. Identity is the number of matching non-gap symbols divided by alignment length. The random-DNA reference in panel d is evaluated by this same alignment procedure. Panels d and f weight identities by reference protein length; panel e averages paired visible-minus-shuffled differences equally across its 32 CDSs. Confidence intervals resample source CDSs, preserving pairs. The full-mask categorical budgets are 32 steps for CPDD and 256 for PhaGen; visible-copy scoring in panel e uses single-pass argmax.

### 3.14 Model-free nucleotide reference ladder

Figure 5a evaluates six reference rules on the same 24 target CDSs, totaling 17,238 nucleotides (5,746 codons). The complete target CDS is masked for model scoring; surrounding sequence remains visible. Reference rules instead use the information specified in Table 2. Targets are oriented in the coding direction using their annotated strand and reading frame. All coding positions, including the first and terminal codons, enter the nucleotide-accuracy denominator. The score is base-by-base agreement with the original DNA, not amino-acid identity or the fraction of exactly recovered codons.

**Table 2:**
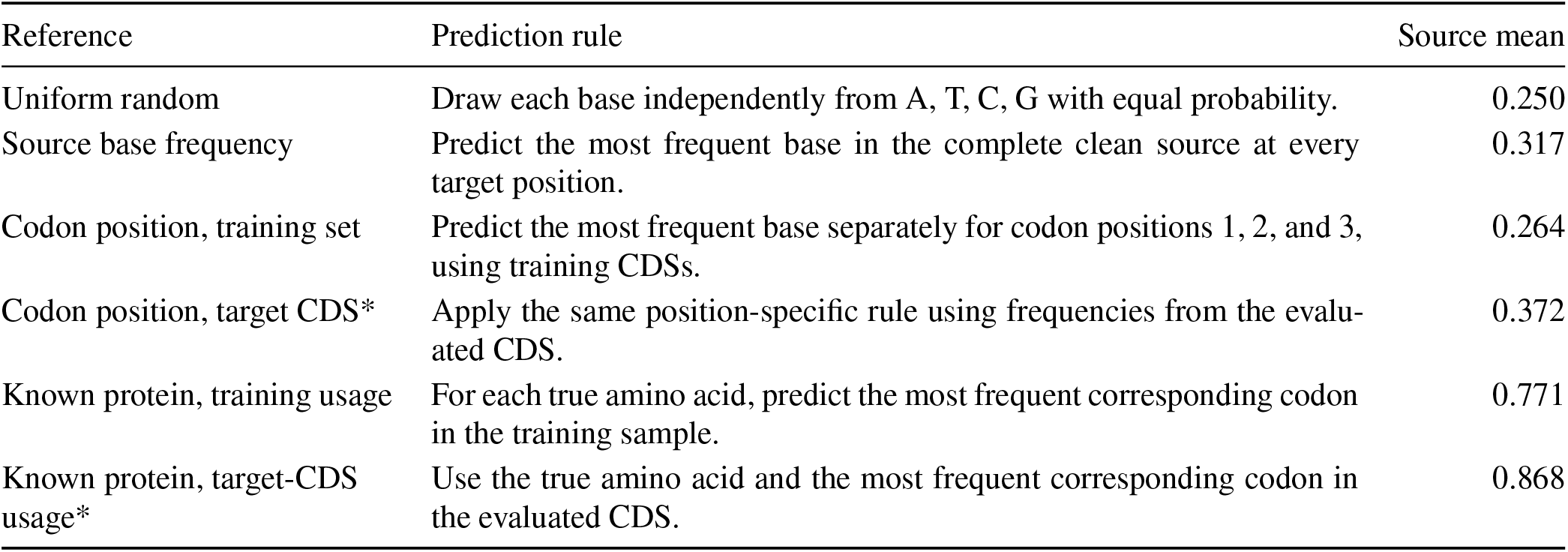
Reference rules used in Figure 5a. All accuracies are unweighted means across the same 24 source CDSs. The uniform rule reports an analytical expectation. Asterisks denote statistics estimated from the evaluated target CDS itself.

| Reference | Prediction rule | Source mean |
|---|---|---|
| Uniform random | Draw each base independently from A, T, C, G with equal probability. | 0.250 |
| Source base frequency | Predict the most frequent base in the complete clean source at every target position. | 0.317 |
| Codon position, training set | Predict the most frequent base separately for codon positions 1, 2, and 3, using training CDSs. | 0.264 |
| Codon position, target CDS* | Apply the same position-specific rule using frequencies from the evalu-ated CDS. | 0.372 |
| Known protein, training usage | For each true amino acid, predict the most frequent corresponding codon in the training sample. | 0.771 |
| Known protein, target-CDS usage* | Use the true amino acid and the most frequent corresponding codon in the evaluated CDS. | 0.868 |

Training frequencies are pooled from 3,000 valid whole CDSs sampled from the eligible training pool after applying the frozen source-exclusion list (random seed 20261105). Empirical counts are normalized without pseudocounts. For the source-base rule, frequencies are counted over the complete source, including the target, and complemented for negative-strand targets so that the distribution and target use the same coding orientation. Codon-position rules choose a base independently at each of the three reading-frame positions and repeat these choices across the CDS; they do not select a complete codon jointly. A training-derived rule can therefore perform below the uniform expectation when its preferred bases differ from those of the evaluation panel.

The protein-conditioned rules use the target’s true amino-acid labels under its annotated genetic code and choose a complete codon by maximum observed frequency. The first codon uses a separately counted START category, and stop codons use a stop category. If a category is absent from the training counts, the rule uses a uniform distribution over the corresponding codons permitted by the target genetic code; argmax ties follow the stored category order. These are references with privileged information: both protein-conditioned rules receive the true protein, the asterisked rules read the hidden target’s statistics, and the source-base rule also includes target bases in its composition estimate. They are not deployable restoration baselines or formal performance ceilings.

For reference rule *r*, the displayed accuracy is

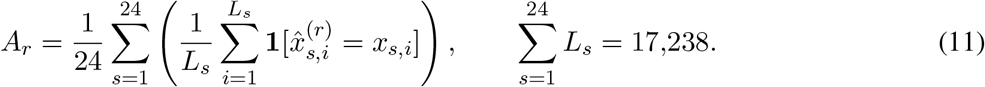

Thus, each CDS first contributes its own correct-base fraction, and these 24 fractions are averaged. For example, the training codon-position rule gives 0.26437 by this procedure; pooling its 4,283 correct calls over 17,238 bases would give 0.24846, a length-weighted statistic that is not plotted. The uniform reference is exactly ¼ in expectation. Model lines use the same source weighting, averaging replicates within each source before averaging across sources, with 95% percentile intervals from 10,000 source-bootstrap resamples. The ladder provides descriptive context, not a formal performance ceiling or an estimate of the fraction of recoverable information captured by a model.

## 4 Discussion

CPDD demonstrates a practical route for converting an AR phage model with conditional memory and depth-wise residual mixing into a bidirectional denoiser. AR initialization reuses sequence features learned during pretraining, while the base-5 Engram avoids discarding every *k*-mer touching a mask. The staged objective also exposes the model to both absent and incorrect visible bases, supporting prediction in inputs that contain a mixture of missing and potentially erroneous sequence.

The scaffold-based evaluations show random-mask restoration well above codon-level references, informative frozen representations, and recovery of visible incorrect bases. Comparisons with a model of the same class provide context for these measurements. Together, the results motivate further development of local refill and editing within an existing sequence scaffold.

Longer-range restoration offers a shared direction for further development of diffusion DNA language models. The controlled evaluations show a stronger contribution from codon-usage statistics as visible context becomes less informative, while copy-bearing training examples support accurate recovery of supervised sequences. The limited transfer observed in held-out copy contexts motivates readout-oriented training and objectives that couple codons across a gene; the relative contributions of training signal, architecture, and decoding remain open. These findings provide a basis for further development of scaffold-conditioned sequence models. The present interface restores fixed-length targets within a preserved scaffold; insertions, deletions, and coordinated whole-genome design remain directions for future development. The basic decoder retains earlier writes, and re-mask repair has not yet produced a net identity gain at the tested budgets. Distinguishing recovery at planted error sites from end-to-end repair will therefore remain important when developing editing applications.

The evaluation panels were screened against documented training ranges but are enriched for *Pseudomonas aeruginosa*-related phages, and distant homology cannot be excluded. Some comparisons use unequal sampling budgets or adapted protocols, as specified in the figure legends, and CPDD is not guaranteed reverse-complement equivariant. The results characterize computational sequence recovery and representation quality; biological activity has not been experimentally evaluated.

The longer-term aim is modular design of phage genomes, with selected coding regions or modules modeled in their surrounding genomic context. Progress towards this aim may benefit from training that rewards context readout, exposure to the model’s own predictions, and codon- or protein-joint objectives that coordinate recovery across a coding region. Evaluating both corrected errors and unintended edits, together with independent functional validation, will be essential for translating these modeling advances into sequence-design applications.

## Supporting information

demo

## Data and code availability

The underlying public sequence collection follows megaDNA [3] and its NCBI, MGV [15], and GPD [16] sources. Numerical figure data and plotting scripts are included with the manuscript source. Code is available on GitHub at https://github.com/dhnihaoya/CreatiPhage-Diffusion-DNA-Lm (Apache-2.0). Model weights are hosted on Hugging Face (CC BY 4.0) at https://huggingface.co/dinghhhhhhhhhhhhhhh/CreatiPhage-145M and https://huggingface.co/dinghhhhhhhhhhhhhhh/CPDD-147M.

## Acknowledgements

We thank the researchers at DeepSeek-AI and Peking University behind Engram, the Kimi team at Moonshot AI behind Attention Residuals, and the teams behind megaDNA, PhaGen, D3LM, LLaDA, DiffuLLaMA, and GIDD for making their research and associated resources publicly available.

## References

[1] Nguyen, E., Poli, M., Faizi, M., Thomas, A., Wornow, M., Birch-Sykes, C., et al. HyenaDNA: Long-Range Genomic Sequence Modeling at Single Nucleotide Resolution. Advances in Neural Information Processing Systems, 36:43177–43201, 2023.

[2] Nguyen, E., Poli, M., Durrant, M. G., Kang, B., Katrekar, D., Li, D. B., et al. Sequence modeling and design from molecular to genome scale with Evo. Science, 386(6723):eado9336, 2024. doi: 10.1126/science.ado9336.

[3] Shao, B. and Yan, J. A long-context language model for deciphering and generating bacteriophage genomes. Nature Communications, 15:9392, 2024. doi: 10.1038/s41467-024-53759-4.

[4] Ji, Y., Zhou, Z., Liu, H., and Davuluri, R. V. DNABERT: pre-trained Bidirectional Encoder Representations from Transformers model for DNA-language in genome. Bioinformatics, 37(15):2112–2120, 2021. doi: 10.1093/bioinformatics/btab083.

[5] Dalla-Torre, H., Gonzalez, L., Mendoza-Revilla, J., Lopez Carranza, N., Grzywaczewski, A. H., Oteri, F., et al. Nucleotide Transformer: building and evaluating robust foundation models for human genomics. Nature Methods, 22:287–297, 2025. doi: 10.1038/s41592-024-02523-z.

[6] Austin, J., Johnson, D. D., Ho, J., Tarlow, D., and van den Berg, R. Structured Denoising Diffusion Models in Discrete State-Spaces. Advances in Neural Information Processing Systems, 34:17981–17993, 2021.

[7] Sahoo, S. S., Arriola, M., Schiff, Y., Gokaslan, A., Marroquin, E. M., Chiu, J. T., et al. Simple and Effective Masked Diffusion Language Models. Advances in Neural Information Processing Systems, 37: 130136–130184, 2024.

[8] Nie, S., Zhu, F., You, Z., Zhang, X., Ou, J., Hu, J., et al. Large Language Diffusion Models. Advances in Neural Information Processing Systems, 38, 2025.

[9] Gong, S., Agarwal, S., Zhang, Y., Ye, J., Zheng, L., Li, M., et al. Scaling Diffusion Language Models via Adaptation from Autoregressive Models. In International Conference on Learning Representations, 2025.

[10] Yang, Z., Liu, H., Cao, C., and Su, B. D3LM: A Discrete DNA Diffusion Language Model for Bidirectional DNA Understanding and Generation. arXiv:2603.01780, 2026. MLGenX 2026 workshop paper.

[11] Hou, Y., Long, W., Su, H., Feng, J., and Zhang, Y. PhaGen: masked diffusion for bacteriophage genome analysis and fast genome-scale generation. Public model implementation, pretrained weights, and project page, 2026. URL https://ai4nucleome.github.io/PhaGen/. Accessed 5 November 2026.

[12] Cheng, X., Tian, R., Zeng, W., Dai, D., Chen, Q., Wang, B., et al. Conditional Memory via Scalable Lookup: A New Axis of Sparsity for Large Language Models. arXiv:2601.07372, 2026. URL https://github.com/deepseek-ai/Engram.

[13] Kimi Team, Chen, G., Zhang, Y., Su, J., Xu, W., Pan, S., et al. Attention Residuals. arXiv:2603.15031, 2026. URL https://github.com/MoonshotAI/Attention-Residuals.

[14] von Rütte, D., Fluri, J., Ding, Y., Orvieto, A., Schölkopf, B., and Hofmann, T. Generalized Interpolating Discrete Diffusion. Proceedings of the 42nd International Conference on Machine Learning, 267:61810–61843, 2025.

[15] Nayfach, S., Páez-Espino, D., Call, L., Low, S. J., Sberro, H., Ivanova, N. N., et al. Metagenomic compendium of 189,680 DNA viruses from the human gut microbiome. Nature Microbiology, 6:960–970, 2021. doi: 10.1038/s41564-021-00928-6.

[16] Camarillo-Guerrero, L. F., Almeida, A., Rangel-Pineros, G., Finn, R. D., and Lawley, T. D. Massive expansion of human gut bacteriophage diversity. Cell, 184(4):1098–1109.e9, 2021. doi: 10.1016/j.cell.2021.01.029.

[17] Camargo, A. P., Roux, S., Schulz, F., Babinski, M., Xu, Y., Hu, B., et al. Identification of mobile genetic elements with geNomad. Nature Biotechnology, 42(8):1303–1312, 2024. doi: 10.1038/s41587-023-01953-y. Online publication 2023.

[18] Lin, Z., Akin, H., Rao, R., Hie, B., Zhu, Z., Lu, W., et al. Evolutionary-scale prediction of atomic-level protein structure with a language model. Science, 379(6637):1123–1130, 2023. doi: 10.1126/science.ade2574.

[19] Hou, Y., Long, W., Su, H., Feng, J., and Zhang, Y. PhaGen: code, evaluation data and results. Zenodo, 2026. URL https://zenodo.org/records/23477083.

[20] Kelsic, E. D., Chung, H., Cohen, N., Park, J., Wang, H. H., and Kishony, R. RNA Structural Determinants of Optimal Codons Revealed by MAGE-Seq. Cell Systems, 3(6):563–571.e6, 2016. doi: 10.1016/j.cels.2016.11.004.

[21] Shang, J., Peng, C., Tang, X., and Sun, Y. PhaVIP: Phage VIrion Protein classification based on chaos game representation and Vision Transformer. Bioinformatics, 39(Supplement_1):i30–i39, 2023. doi: 10.1093/bioinformatics/btad229.

[22] Ondov, B. D., Treangen, T. J., Melsted, P., Mallonee, A. B., Bergman, N. H., Koren, S., et al. Mash: fast genome and metagenome distance estimation using MinHash. Genome Biology, 17:132, 2016. doi: 10.1186/s13059-016-0997-x.

[23] Hyatt, D., Chen, G.-L., LoCascio, P. F., Land, M. L., Larimer, F. W., and Hauser, L. J. Prodigal: prokaryotic gene recognition and translation initiation site identification. BMC Bioinformatics, 11:119, 2010. doi: 10.1186/1471-2105-11-119.

[24] Steinegger, M. and Söding, J. MMseqs2 enables sensitive protein sequence searching for the analysis of massive data sets. Nature Biotechnology, 35(11):1026–1028, 2017. doi: 10.1038/nbt.3988.

[25] Terzian, P., Olo Ndela, E., Galiez, C., Lossouarn, J., Pérez Bucio, R. E., Mom, R., et al. PHROG: families of prokaryotic virus proteins clustered using remote homology. NAR Genomics and Bioinformatics, 3(3): qab067, 2021. doi: 10.1093/nargab/lqab067.

